# Genome-wide identification and transcriptional analyses of the R2R3-MYB gene family in wheat

**DOI:** 10.1101/2020.10.17.343632

**Authors:** Chunsheng Xiao, Yong Jia, Calum Watt, Wenshuai Chen, Yujuan Zhang, Chengdao Li

**Affiliations:** Triticeae Research Institute, Sichuan Agricultural University, Chengdu 611130, China; State Agricultural Biotechnology Centre (SABC), College of Science, Health, Engineering and Education, Murdoch University, WA, 6150, Australia; Western Crop Genetic Alliance, Murdoch University, WA, 6150, Australia; Department of Primary Industry and Regional Development, Government of Western Australia, South Perth, WA, 6155, Australia

**Author notes:** These authors have contributed equally to the study (Chunsheng Xiao, Yong Jia). Correspondence: Prof. Chengdao Li.

**Keywords:** gene duplication, R2R3-MYB, wheat

## Abstract

MYB transcription factors (TFs) represents one of the largest TF families in plants. In this study, we performed genome-wide MYB-domain screening and identified a total of 997 MYBs in wheat *(Triticum aestivum),* among which 445 were 2-domain MYBs (R2R3-MYBs) that were clustered into 15 subgroups with varied conservation profiles. Homologous genes were highly conserved across the three subgenomes, with minor variations contributed by segmental duplications. Tandem and proximal gene duplications have contributed significantly to the expansion of the wheat *Myb* gene family. Furthermore, comprehensive transcriptome profiling of *R2R3-Myb* genes in 61 different tissue and time point samples revealed a clear pattern of temporal and spatial variations within six expression groups. The comprehensive genomic and transcriptional analyses provided valuable insights into the evolution and biological functions of *R2R3-Myb* genes in wheat. They would serve as a useful guide to further investigate the potential agronomic traits controlled by this large TF family.

## 1. Introduction

The myeloblastosis (MYB) transcription factors (TFs) constitute one of the most prominent TFs families in plants. Gene expression regulation by MYBs plays a critical role in plant development and environmental adaption [1,2]. MYBs typically feature the presence of 1~4 repeats of the DNA-binding domain at the N-terminal [3]. The tertiary structure of each repeat encodes three α-helices, with the second and third helices forming a helix-turn-helix (HTH) structure that binds to specific target sequence motifs. The third helix determines the recognition helix that makes direct contact with the major groove of DNA [4]. The C-terminus of MYBs represents the most diverse region and functions as the trans-acting domain (TAD). The diversity of TAD in MYBs is the driver of their broad functional activities [5].

A total of 3,485 MYBs and 2,754 MYB-related sequences are available in the plant TF database (http://planttfdb.cbi.edu.cn/). Depending on the number of DNA-binding domain repeats, MYBs have been classified into four classes:1R–MYB or MYB-related proteins (single repeat), R2R3-MYB (two repeats), 3R–MYB (three repeats), and 4R–MYB (four repeats) [1]. After the first identification of plant *Myb* gene COLORED1 (C1) that involved in anthocyanin biosynthesis in maize [6], many *Myb* genes have been identified in various plant species, such as *Arabidopsis* (405) [1], rice (155) [7], maize (158) [8], soybean (252) [9] and watermelon (162) [10]. Most plant Myb genes encode proteins of the R2R3-MYB class [11,12], which are thought to have evolved from *3R-Myb* genes, by the loss of the sequences encoding the R1 repeat and subsequent expansion of the gene family [13,14]. Many plant R2R3-MYB proteins have been functionally characterized and were found to regulate diverse plant-specific processes, including primary and secondary metabolic pathways [15], plant development [16], signal transduction [17], and stress responses [18].

Wheat *(Triticum aestivum)* is one of the most important cereals, supplying ~ 20% of the calorific need (Food and Agriculture Organization of the United Nations, 2013). At least 73 *Myb* genes have been identified in wheat and perform diverse functions in plant-specific processes [19]. For example, *TaPL1* acts as a positive regulator of anthocyanin biosynthesis in wheat coleoptiles [20]. The ectopic expression of wheat *TaMyb18* in rice was reported to have a dose-dependent effect on the regulation of leaf rolling traits [21]. In addition, *TaMyb30* and *TaMyb31* play critical roles in drought-response [22,23]. *TaRIM1* positively regulates resistance response to Rhizoctonia cereals infection by modulating the expression of a range of defense genes and represents a candidate gene for sharp eyespot resistance in wheat [24].

In this study, we performed a genome-wide identification of 445 R2R3-MYB TFs in wheat and carried out phylogeny, gene structure, gene duplication, and synteny analyses. Comprehensive transcriptome profiling based on RNAseq was performed for the identified *R2R3-Myb* genes. This study improves our understanding of the wheat *R2R3-Myb* gene family and represents a valuable resource for future gene cloning and functional analysis studies of MYB TFs in wheat.

## 2. Results

### 2.1. Identification of R2R3-Myb genes in wheat and gene duplication, synteny analyses

In this study, MYB domains were identified by screening against the Pfam profile PF00249 (Myb-like DNA-binding domain). According to the MYB domain annotation in *Arabidopsis,* a typical MYB domain is ~47 amino acids. A preliminary BLASTP search was performed using the MYB domain sequences of known *Arabidopsis* MYB proteins as queries in the wheat genome. To determine the accurate number of *R2R3-Myb* genes in the wheat, i.e., those genes with two MYB domains, we took the potential presence of partial MYB domain fragment into consideration and counted the MYB domains based on different domain-length thresholds. The number of genes classified into different categories varied when thresholds ≥ 20aa and ≥ 25aa in comparison with that at ≥ 30aa, ≥ 35aa, and ≥ 40aa, suggesting the presence of partial MYB domains (**Table 1**). The numbers of genes identified as 3-, 4-, 5- and 6-domains are consistent at ≥ 30aa, ≥ 35aa, and ≥ 40aa. Thus, we defined that a threshold of domain-length above 30aa denoted possible R2R3-MYB, which resulted in the identification of 997 *Myb* genes, including 454 *R2R3-Myb* genes in the wheat genome.

**Table 1.**
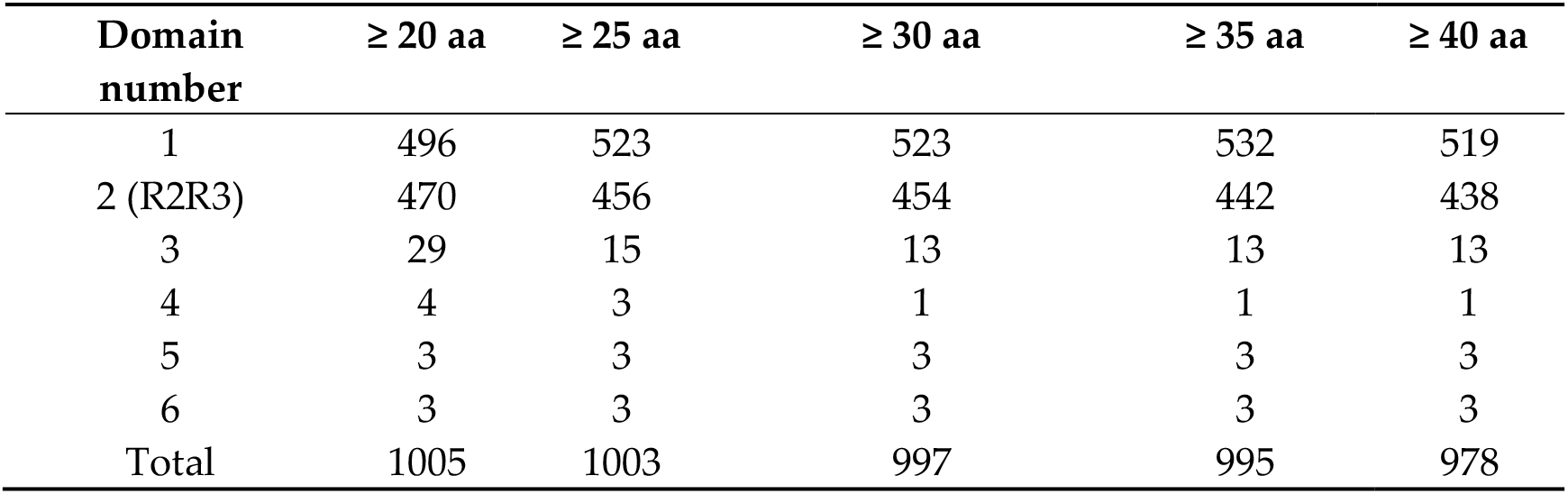
MYB domain screening of wheat genome. Domain number was counted based on a different threshold (≥ 20, ≥ 25, ≥ 30, ≥ 35, ≥ 40 aa). (See Supplementary **Table S1** for detailed gene ID lists)

| Domain number | $\geq 20$ aa | $\geq 25$ aa | $\geq 30$ aa | $\geq 35$ aa | $\geq 40$ aa |
| --- | --- | --- | --- | --- | --- |
| 1 | 496 | 523 | 523 | 532 | 519 |
| 2 (R2R3) | 470 | 456 | 454 | 442 | 438 |
| 3 | 29 | 15 | 13 | 13 | 13 |
| 4 | 4 | 3 | 1 | 1 | 1 |
| 5 | 3 | 3 | 3 | 3 | 3 |
| 6 | 3 | 3 | 3 | 3 | 3 |
| Total | 1005 | 1003 | 997 | 995 | 978 |

Of the 997 *Myb* genes in the wheat genome, ~ 80.6 % (804) were found in conserved collinear regions and were thus determined as whole-genome duplication (WGD)/segmental duplications. In addition, ~ 15.7% were identified as tandem duplications (135) and proximal duplications (22). Only ~ 3.1 % of genes (31) were considered as dispersed duplications. These results suggest that the majority of *Myb* genes were conserved across the A, B, and D sub-genomes. Tandem duplication and proximal duplications have, however, contributed substantially to the expansion of the wheat *Myb* gene family. A similar pattern was observed when 454 *R2R3-Myb* genes were examined individually, which contain 365 (~ 80.4%), 63 (~ 13.9%), 7 (~ 1.5%), and 17 (~ 3.7%) WGD/segmental, tandem, proximal and dispersed duplications, respectively.

In total, 332, 327, and 333 *Myb* genes were identified in A, B, and D sub-genomes, respectively (**Figure 1a-c**). Each sub-genome displayed a comparable *Myb* genetic location and synteny profile, coinciding with the gene duplication pattern results, which showed the majority of *Myb* genes were located in conserved collinear regions. Despite this, several significant variations of *Myb* collinearity in different sub-genomes were identified. For example, a clear *Myb* collinearity cluster between the chromosome tip of 7A and the chromosome end of 4A were identified, which were not present in B and D sub-genomes (**Figure 1a-c**). This may reflect a sub-genome specific segmental duplication event. Moreover, unique inter-chromosome collinearity was identified on chromosome 7B (**Figure 1b**). For all sub-genomes, we observed a clear domain-number specific clustering pattern for the 1-domain, 2-domain (R2R3-Myb), and 3-domain genes. The majority of the identified collinearity lines were also domain-number specific, suggesting general conservation of domain number in different types of *Myb* genes. However, a few gene clusters represented by collinear genes with variable amounts of MYB domains could also be observed (**Figure 1a-c**), suggesting the occurrence of domain gain or loss events during gene family evolution.

**Figure 1.**
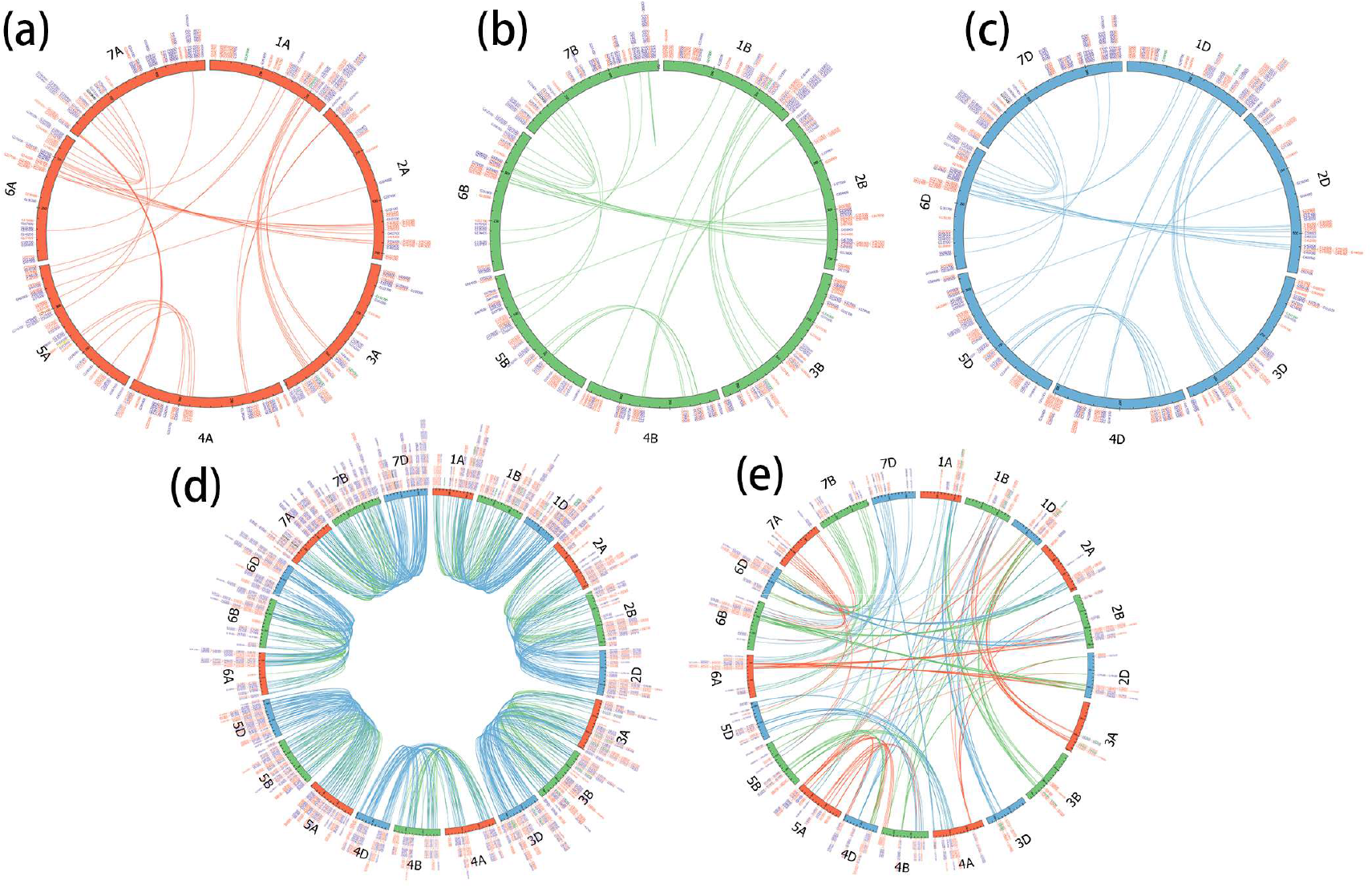
Synteny analyses of MYB domain-containing genes in wheat. *Myb* gene collinearity was identified using MCScanX tool. Collinearity within each subgenome A (**a**: red), B (**b**: green), D (c: blue) was displayed, followed by inter-sub-genome collinearity by chromosome (**d**) and inter-sub-genome collinearity (**e**). For **(a)**, **(b),** and **(c)**, all MYB-domain genes identified by the ≥ 30 aa criteria were labeled to display the distribution of *Myb* genes across the wheat genome. For **(d)** and **(e)**, only those identified as collinear genes were labeled. Each *Myb* gene pair was linked by curved lines, colored according to the chromosome color of the second *Myb* gene. Standard gene ID “TraesCS[1–7] [A/B/D]02Gxxxxxx” were shortened as “Gxxxxxx” and colored according to the number of MYB domains (1: purple; 2: red, i.e. *R2R3-Myb;* 3: green; 4: yellow; 5: orange; 6: black).

In addition to the intra-sub-genome collinearity, inter-sub-genome *Myb* collinearity was displayed in **Figure 1a** and **1e**. Results revealed the general conservation of *Myb* genes across A, B, and D sub-genomes for chromosomes 1-7 (**Figure 1d**). In contrast, the interchromosome collinearity reflected the potential inter-chromosome segmental duplication events and displayed a much less conserved pattern. These identified patterns of interchromosome collinearity provide valuable insight into the complex expansion and evolutionary history of the wheat *Myb* gene family.

### 2.2 Classification of wheat R2R3-MYBs

To gain insight into the evolutionary relationship of R2R3-MYBs in wheat, we developed a Neighbor-Joining (NJ) phylogeny (**Figure 2**) according to protein sequence similarity. We separated the 454 wheat R2R3-MYBs into 15 subgroups (designated WS1-WS15) (**Figure 2**). WS2, WS4, WS5, and WS7 contained no more than six R2R3-MYBs. It was likely because the MYB domain was relatively short, and that members within a subgroup were highly conserved, with some unique sequence characters. WS6 and WS10 were the two largest subgroups, and each had 62 members. We also compared the subgroups of *Arabidopsis.* The clustering pattern of *Arabidopsis* MYBs is highly consistent with the previous classification [2], indicating the reliability of this phylogenetic approach.

**Figure 2.**
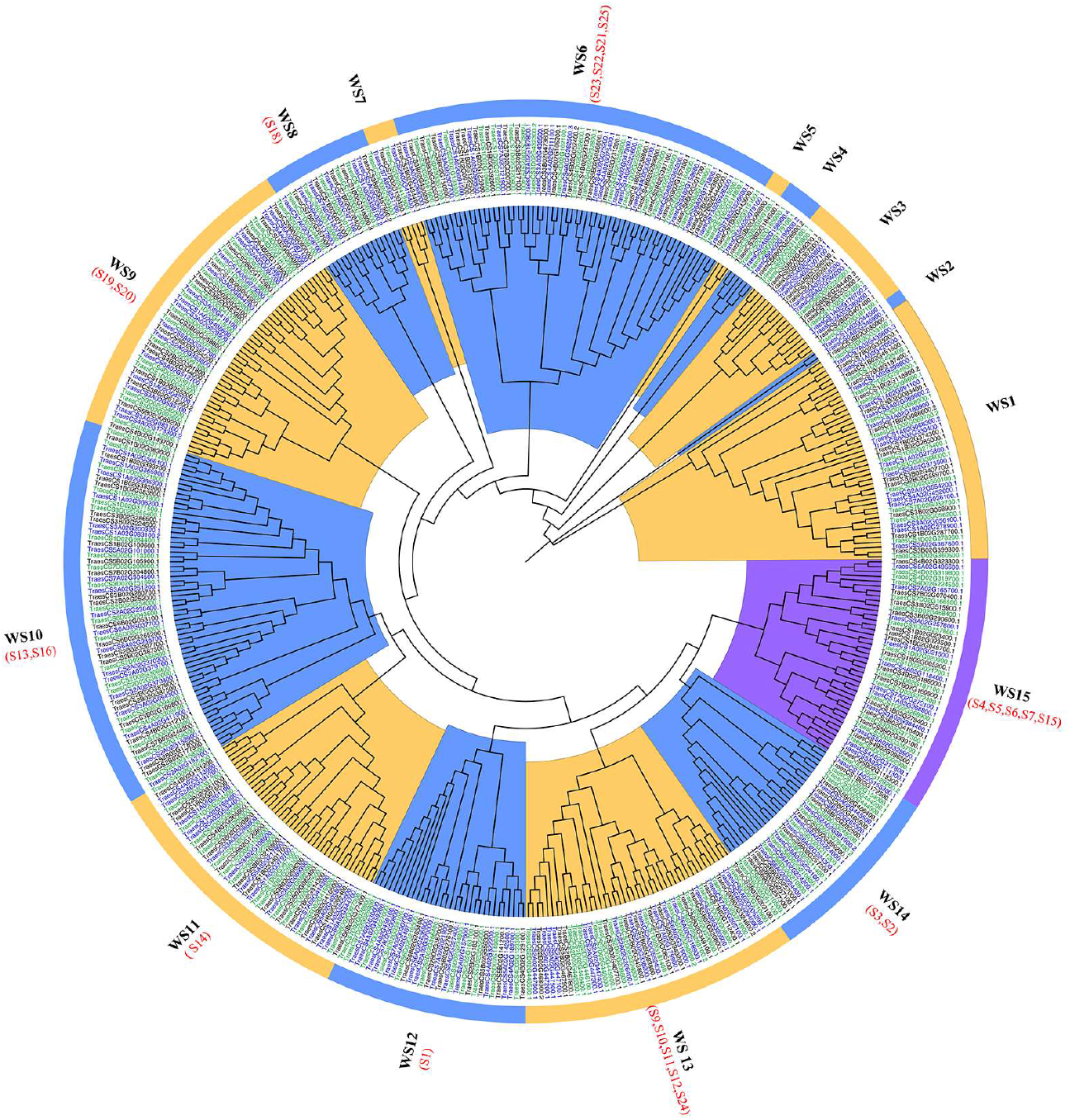
Phylogenetic tree of wheat MYB TFs. The Neighbor-Joining phylogeny was developed based on amino acid sequence alignment of the MYB domains of wheat R2R3-MYBs (gene ID in blue, black, and green for subgenomes A, B, and D, respectively). Wheat Subgroups (WS1-WS15) were labeled by camber lines. Subgroup labels of *A. thaliana* R2R3-MYBs were marked by red (S1-S25).

Most of the R2R3-MYB subgroups in *Arabidopsis* were present in the wheat genomes (**Figure 2**), with the expectation of S8 and S17. However, wheat R2R3-MYBs in WS1~WS5 and WS7 cannot be assigned to any specific subgroup using *Arabidopsis* MYBs as references. These R2R3-MYBs may represent wheat specific subgroups due to the remote relationship between the two species.

### 2.3 Gene structure analyses

Gene structural elements, including 5’-UTR, exon, intron, and 3’-UTR contain essential information to trace gene evolution. Gene fundamental data for each wheat *R2R3-Myb* subgroup were analyzed based on the wheat genome annotation database and visualized (**Figure 3**). Overall, the *R2R3-Myb* gene structure displayed a clear distinction across different subgroups, supporting a common evolutionary origin for individual subgroups. It also indicates that the R2R3-MYB subgroups had diverged in the distant past. However, the number of introns in wheat *R2R3-Myb* genes appeared to be limited. The majority (55.28%) of the 454 *R2R3-Myb* genes had no more than two introns. A large number (44.05%) of the R2R3-type domain-containing proteins had a conserved splicing pattern with three exons and two introns, while 51 (11.23%) *R2R3-Myb* genes belonging to WS2, WS4, WS9, and WS10 were intronless (**Figure 3**). WS2, WS7, WS4, WS8, WS3, WS11, and WS12 were relatively more conserved, including the length and numbers of exons, suggesting a potentially conserved biological function. In contrast, the structures of the genes from other subgroups, especially WS1 and WS9, appeared more divergent in the number of exons. In addition, four and seven R2R3-MYB genes in WS4 and WS1 contained more than 11 exons, which was not found in *Arabidopsis R2R3-Myb* genes. We also observed significant intra-subgroup differences in terms of the presence, loss, and length-variation of 5’-UTRs, 3’-UTRs, which were present in most subgroups. These genomic differences may have contributed to the potential transcriptional divergence for the *R2R3-Myb* gene members within each subgroup.

**Figure 3.**
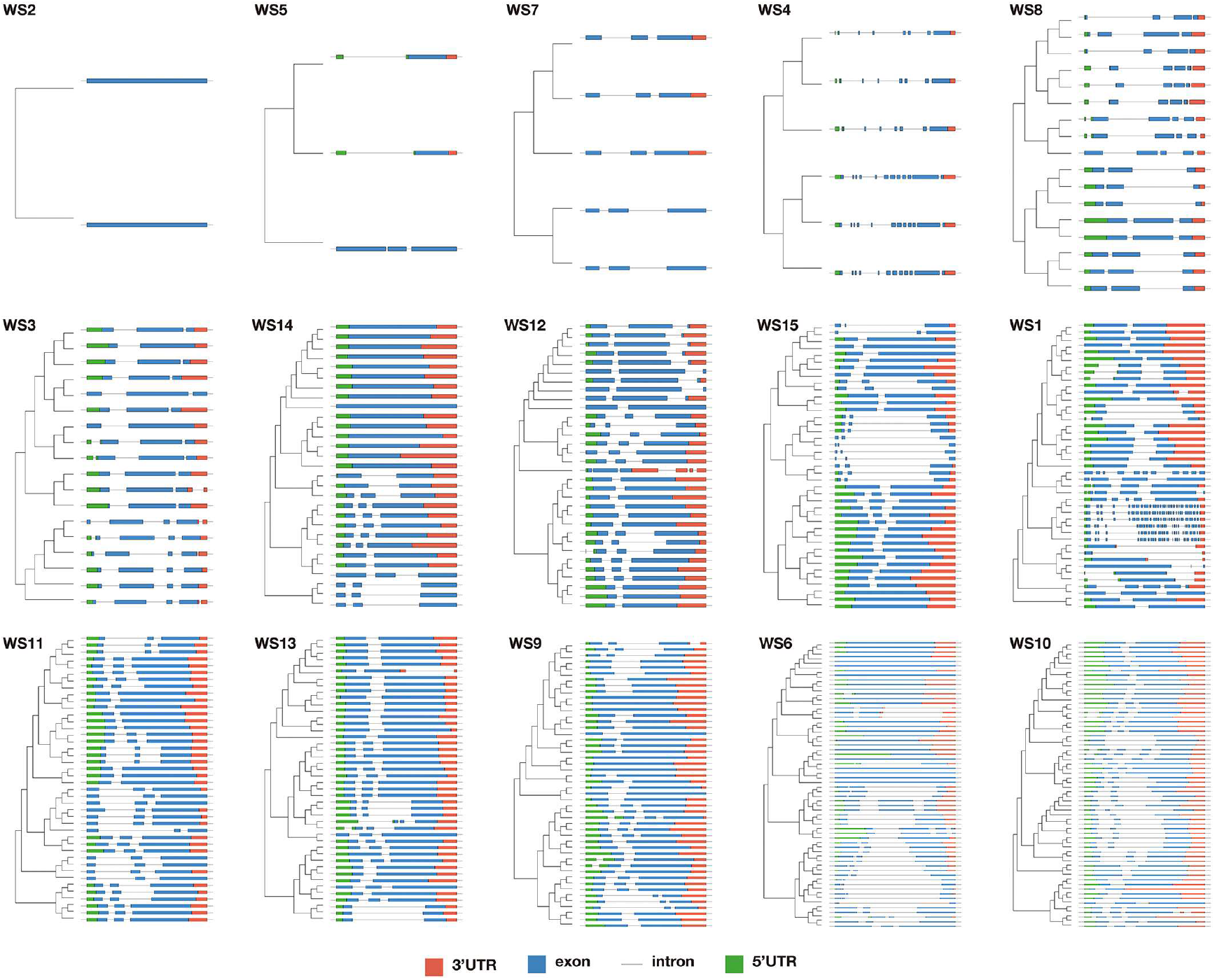
Gene structure of wheat *MYB* genes. The subgroup was labeled according to phylogeny classification in this study. Gene structure was based on the wheat genome annotation. 3’-UTR (red), 5’-UTR (green), exon (blue), and intron (line) were displayed in schematic diagrams.

### 2.4 Transcriptional profiling

The expression patterns of *R2R3-Myb* genes were characterized using the 61 RNA-seq data for different wheat growth stages or tissues (**Table S2; Figure 4**). Eight groups (G1-G8) of R2R3-MYB TFs with significantly altered expression levels were detected (**Figure 4**). The transcriptional data for different groups displayed a clear pattern of temporal and spatial variations. Significant transcriptional differences were also prevalent among individual genes within each group, reflecting an extensive degree of transcriptional divergence among the wheat *R2R3-Myb* gene family. According to transcriptional profiling, the majority of genes in G1, G2, and G8 were highly expressed in most detected tissues. The majority of genes in G2 and those G1 genes were highly expressed in the root, while they had no expression in the endosperm. We also identified 163 genes in G3 that barely expressed in the majority of tissues. Root tissue displayed the detectable transcription for around 2/3 of the genes identified, especially in G4, all of the genes had a different level of transcript accumulation in all nine root samples. In addition, genes in G5, G6, and G7 also showed significant tissue-specific expression. For example, 41 genes in G5 were mainly expressed in developing embryo and revealed significantly altered expression during development. Nine genes in G7 displayed high expression levels in developing seed (10DPA-30DPA), including grain, endosperm, starch, aleurone, and transfer cells. G6 consisted of 13 genes belonging to WS8, WS9 had the highest transcription in anther and spike during anthesis. Highly expressed genes in other flower-related tissues (ovary, stigma, pistils, stamen, peduncle, and rachis) could also be identified in G2 and G8.

**Figure 4.**
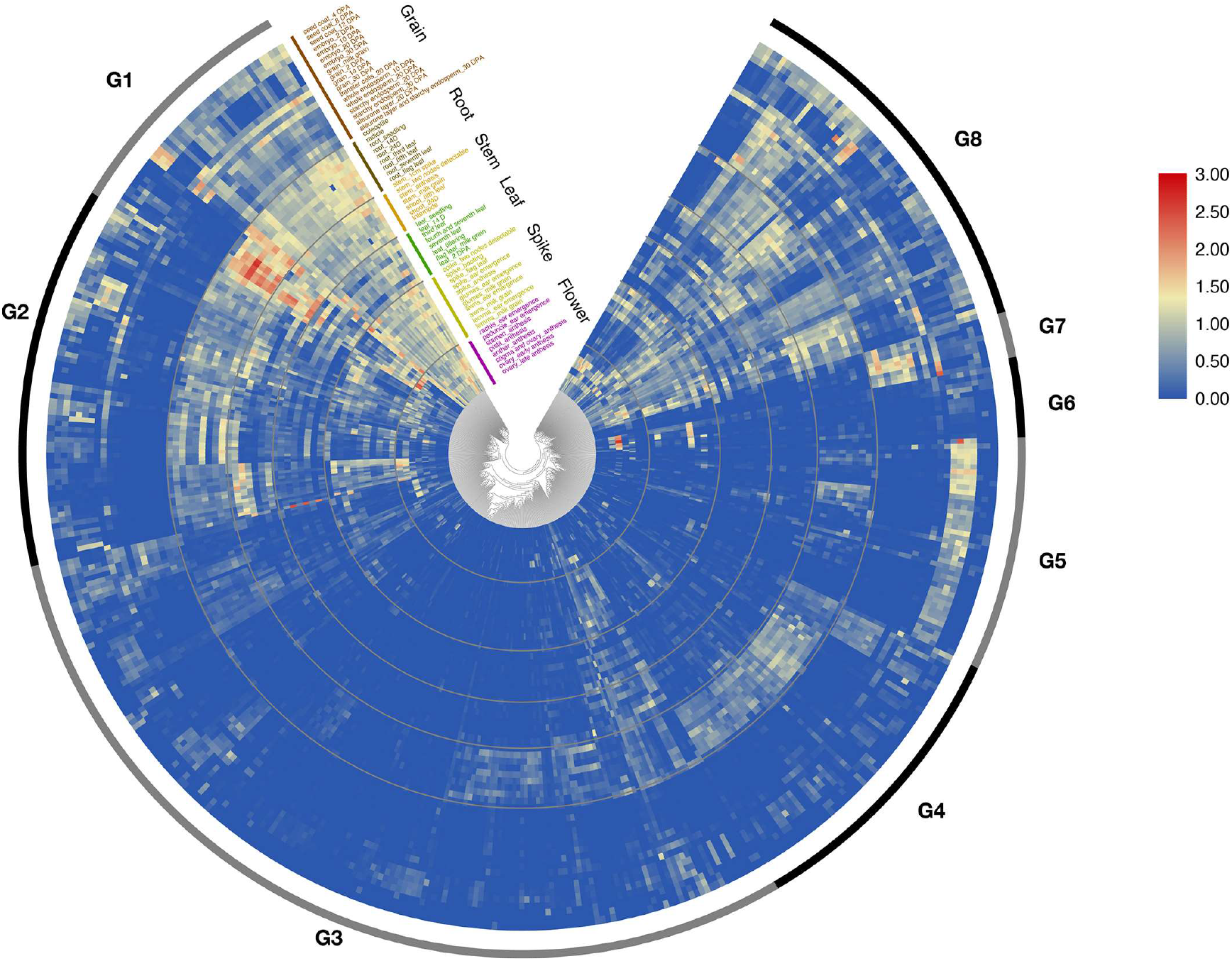
Transcriptional profiling of wheat *R2R3-Myb* genes in cv. Chinese Spring. RNAseq data for 61 different samples were downloaded from NCBI database. *Day(D); Days post-anthesis (DPA).

## 3. Discussion

The MYB TF gene family is one of the largest TF families in plants and has been identified to be involved in regulating numerous plant physiological and biochemical processes. Within the MYB TFs family, the 2-domain MYB type, i.e., R2R3-MYB, accounts for a large proportion and has been the main focus in previous studies. Phylogenetic analyses of R2R3-MYBs have been systematically studied in *Arabidopsis* [1,2] and many other species [7–9], which indicated that its members perform diverse functions in plant-specific processes. However, knowledge of R2R3-MYBs in wheat are had been limited for a long time due to its complex genome composition and lack of reference genome data [19]. The increasing availability of wheat genome sequences provides more possibility for the study of R2R3-MYBs in wheat [25].

In the present study, we performed a genome-wide investigation of the *MYB* gene family in wheat and identified a total of 997 *Myb* genes, including 454 putative wheat *R2R3-Myb* genes. The number of wheat R2R3-MYBs is much higher than an earlier report in wheat. This expansion of MYB TF family in wheat is still very significant, even considering its triplicate subgenomes. Our results showed that gene duplications have significantly contributed to the MYB TFs family expansion. Gene duplication contributes significantly to the expansion of the gene family in the plant kingdom, which leads to the diversification and evolution of genes [26,27]. Depending on the different originating mechanisms, gene duplication can be divided into four categories: whole-genome duplication (WGD) or segmental duplication, tandem duplication, proximal duplication, and dispersed duplication [28]. Our results showed that 365 (~ 80.4%) wheat *Myb* genes were identified as WGD or segmental duplications, whereas only a small proportion (3.9%) of wheat *Myb* genes were identified as dispersed duplications. This observation is surprising, considering that the majority of the wheat genome is comprised of transposable elements [25]. It indicates that wheat *Myb* genes were highly conserved across the three subgenomes. However, we also observed a significant number of inter-subgenome syntenic regions (**Figure 1e**), which have resulted from segmental duplication events. A small amount of subgenome-specific segmental duplications has also been observed in our analyses. Combining domain number and gene synteny analyses revealed the occurrence of domain number changes during gene duplications. Interestingly, in addition to the normal 1-4 domain MYBs, we also identified 1, 3, and 3 MYBs that contain 4-, 5- and 6-MYB domains, respectively. In particular, the 5- and 6-domain *Myb* genes were conserved in all of the A, B, and D sub-genomes, suggesting they evolved early in *Triticeae* genomes. Further investigation is needed to illustrate the possible biological function of these novel genes.

In this study, we divided R2R3-MYBs into 15 subgroups (WS1-WS15), while 25 subgroups (S1-S25) were identified in *Arabidopsis* [2]. The unique subgroup of wheat and *Arabidopsis* reflecting the species-specific evolutionary history of the MYB family or could also be due to the domain-based phylogeny analyses in this study. Indeed, this is not surprising considering the remote distance homology between the two species. Also, the presence of partial MYB domains and the potential domain gain or loss during gene duplications might have contributed to the formation of additional wheat subgroups. It would be interesting for future studies to investigate whether the gene expansion and the formation of unique subgroups have contributed to wheat’s adaptation to broad environment conditions.

Gene structure analysis revealed that the majority of the genes in the same subgroup generally exhibited the same intron pattern, and the position of the intron was also highly conserved, which was consistent with a previous report [9]. Besides, the most common pattern of genes structures and the proportion of intronless *R2R3-Myb* genes in wheat were coincident with previous studies in *Arabidopsis*, rice, and maize [7,29]. This finding proves the reliability of our phylogenetic analysis and constitutes an independent criterion for *Myb* gene family classification.

MYB TFs can activate or inhibit downstream target genes, thus regulating the growth and development of plants [30,31]. Therefore, we used RNAseq data to quantify the expression of *R2R3-Myb* genes in different tissues (**Figure 4**) and combined with a phylogenetic tree to explore the putative functions of the wheat MYB proteins based on their homology with particular functional clades of *Arabidopsis* MYBs (**Figure 2**). The result revealed clear transcriptional divergence within different expression groups, coinciding with previous reports that transcriptional variation is common after gene duplication [32]. Some groups displayed a tissue-specific and developmental regulated transcriptional pattern, which may be related to their diverse biological functions. In G6, three homologs (TraesCS5A02G087100, TraesCS5B02G093000, and TraesCS5D02G099100) of *TaMyb79* belonging to WS9 significantly expressed in anther and the spike during anthesis. *TaMyb79* was an orthologous gene of *AtMyb78* and *AtMyb108,* which function in facilitating proper anther development in *Arabidopsis* [33], indicating that *TaMyb79* might have an essential role in the process of flower development. In addition, TraesCS1A02G406600 and TraesCS1D02G415000 as orthologous genes of *AtMyb21, AtMyb24, AtMyb57* in S19, also had the highest expression level in anther, coinciding with the roles of S19 in stamen development [26]. Four *R2R3-Myb* genes (TraesCS3D02G257800, TraesCS7D02G166500, TraesCS7A02G165700, TraesCS4D02G224500) in WS15 had close phylogenetic relationship with S5S6S15 that were annotated to be responsible for flavonoid and anthocyanins biosynthesis [26]. Those genes were gathered in G5 and were highly expressed in the embryo (**Figure 4**), signifying they were likely to affect seed dormancy due to their predicted function in flavonoid and anthocyanin production [34]. Twelve *R2R3-Myb* genes in WS13 were gathered in G4, where showed high expression in root. One of them was *TaMyb74* (TraesCS2D02G209600), and its expression pattern corresponded to reported studies and its function in the regulation of cuticle biosynthesis [35]. Those genes that came from the same subgroup had similar expression patterns, demonstrating they may play similar roles in regulation.

## 4. Materials and Methods

### 4.1. Sequence retrieval and MYB-domain scanning

The annotated amino acid sequences for the bread wheat genome were downloaded from EnsemblPlants database (https://plants.ensembl.org/index.html). The MYB domain profile was downloaded from the Pfam database (https://pfam.xfam.org/). MYB domain screening was performed using hmmscan tool. R2R3-MYB protein sequences of *Arabidopsis* were retrieved from EnsemblPlants according to gene ID reported in the literature [2].

### 4.2. Sequence alignment and phylogeny

Based on the MYB domain scanning results, sequence fragments for the MYB domains of each target gene were retrieved using in-house Python script. Sequence alignment of the MYB domain sequences for wheat was performed using MUSCLE [36] program (8 iterations). Neighbor-Joining phylogeny was developed using MEGA7.0 [37] with the p-distance substitution model and was validated by bootstrapping for 1000 times. The obtained tree was annotated in FigTree (v1.4.3).

### 4.3. Synteny analysis

Gene duplication and synteny analysis of *Myb* genes in wheat were performed using the MCScanX tool [38]. The protein sequences and gene position data for the A, B, and D sub-genomes of wheat were extracted and treated as separate genomes. All-vs-all blast was performed using NCBI-Blast standalone tool with an *E-value* threshold at 1e-10. The blast results and gene position files were used as inputs for MCScanX. The obtained synteny information was annotated on the genetic map generated using the CIRCOS software [39].

### 4.4. Gene structure analysis

The structural information (5’-UTR, exon, intron and 3’-UTR) for target *Myb* genes were extracted from the gff3 file of the wheat genome (EnsemblPlants release 47) and further modified in format. Gene structure and the phylogeny for each classified MYB subgroup were displayed using the ‘ggtree’ R-software package [40] with in-house scripts.

### 4.5. Expression profiles

We selected RNA-seq data files with tissues or organ information at different wheat developmental stages from Chinese Spring (CS) RNA-Seq data files currently available in the SRA database (http://www.ncbi.nlm.nih.gov/sra). Sixty-one samples included six parts (**Grain**: seed coat, embryo, grain, transfer cells, whole endosperm, starchy endosperm, aleurone layer, aleurone layer, and starchy endosperm; **Root**: coleoptile, radicle, root; **Stem**: stem, shoot, internode; **Leaf**: leaf; **Spike**: spike, glumes, awns, lemma; **Flower**: rachis, peduncle, stamen, pistil, anther, stigma and ovary, ovary). The transcriptional results were visualized in a heatmap using the ‘ggtree’ R-software package [40] and a calculation with log2(TPM+1).

## Supplementary Materials

Table S1. List of identified MYB TFs in wheat.

Table S2. The relative expression of 445 *R2R3-Myb* genes in 61 RNAseq samples.

## Author Contributions

Conceptualization, C. L.; methodology, Y.J.; software, C.W.; validation, Y.J., and Y.Z.; formal analysis, Y.J.; investigation, C.X.; resources, C.X.; writing—original draft preparation, C.X.; review and editing, C.L.; visualization, W.C.; supervision, C.L.; project administration, C.X. All authors have read and agreed to the published version of the manuscript.

## Acknowledgments

We acknowledge the plant research community for sharing the relevant genomic data and making it publicly available.

## Abbreviations

CS: Chinese Spring
D: Days
DPA: Days post-anthesis
HAI: Hours after imbibition
MYB: myeloblastosis
NJ: Neighbor-Joining
TAD: *trans*-acting domain
TFs: Transcription factors
TPM: Transcripts Per Kilobase Million
WGD: Whole-genome duplication
WS: Wheat subgroups

## References

[1] Dubos, C.; Stracke, R.; Grotewold, E.; Weisshaar, B.; Martin, C.; Lepiniec, L. MYB transcription factors in *Arabidopsis*. Trends in Plant Science 2010, 15, 573–581, doi:10.1016/j.tplants.2010.06.005.

[2] Millard, P.S.; Kragelund, B.B.; Burow, M. R2R3 MYB Transcription Factors - Functions outside the DNA-Binding Domain. Trends in Plant Science 2019, 24, 934–946, doi:10.1016/j.tplants.2019.07.003.

[3] Sakura, H.; Kanei-Ishii, C.; Nagase, T.; Nakagoshi, H.; Gonda, T.J.; Ishii, S. Delineation of Three Functional Domains of the Transcriptional Activator Encoded by the C-myb Protooncogene. Proc Natl Acad Sci U S A, 1989, 86, 5758–62, doi:10.1073/pnas.86.15.5758.

[4] Ogata, K.; Morikawa, S.; Nakamura, H.; Sekikawa, A.; Inoue, T.; Kanai, H.; Sarai, A.; Ishii, S.; Nishimura, Y. Solution structure of a specific DNA complex of the Myb DNA-binding domain with cooperative recognition helices. Cell 1994, 79, 639–648, doi:10.1016/0092-8674(94)90549-5.

[5] Du, H.; Tang, XF.; Liu, L.; Yang, W.J.; Wu, Y.M. Huang, Y.B.; Tang, Y.X. Cloning and Functional Identification of Two MYB Transcription Factors GmMYBJ6 and GmMYBJ7 in Soybean. Acta Agronomica Sinica 2008, 34(7), 1179–1187, doi:10.1016/S1875-2780(08)60042-5.

[6] Paz-Ares, J.; Ghosal, D.; Wienand, U.; Peterson, P.A.; Saedler, H. The regulatory C1 locus of Zea mays encodes a protein with homology to MYB-related proto-oncogene products and with structural similarities to transcriptional activators. Embo Journal The EMBO J 1988, 6, 3553–3558, doi:10.1002/j.1460-2075.1987.tb02684.x.

[7] Katiyar, A.; Smita, S.; Lenka, S.; Rajwanshi, R.; Chinnusamy, V.; Bansal, K.C. Genome-wide classification and expression analysis of the MYB transcription factor families in rice and *Arabidopsis*. BMC Genomics 2012, 13, 1–19, doi:10.1186/1471-2164-13-544.

[8] Du, H.; Feng, B.R.; Yang, S.S.; Huang, Y.B.; Tang, Y.X. The R2R3-MYB transcription factor gene family in maize. PLoS One 2012, 7, e37463–e37463, doi:10.1371/journal.pone.0037463.

[9] Du, H.; Yang, S.S.; Liang, Z.; Feng, B.R.; Liu, L.; Huang, Y.B.; Tang, Y.X. Genome-wide analysis of the MYB transcription factor superfamily in soybean. BMC Plant Biol 2012, 12, 106, doi:10.1186/1471-2229-12-106.

[10] Xu, Q.; He, J.; Dong, J.; Hou, X.; Zhang, X. Genomic Survey and Expression Profiling of the MYB Gene Family in Watermelon. Horticultural Plant Journal 2018, 4, 1–15, doi:10.1016/j.hpj.2017.12.001.

[11] Martin, C.; Paz-Ares, J. MYB transcription factors in plants. Trends in Genetics 1997, 13, 67–73, doi:10.1016/S0168-9525(96)10049-4.

[12] Du, H.; Liang, Z.; Zhao, S.; Nan, M.G.; Tran, L.S.; Lu, K.; Huang, Y.B.; Li, J.N. The Evolutionary History of R2R3-MYB Proteins Across 50 Eukaryotes: New Insights Into Subfamily Classification and Expansion. Sci Rep 2015, 5, 11037, doi:10.1038/srep11037.

[13] Rosinski, J.A.; Atchley, W.R. Molecular Evolution of the Myb Family of Transcription Factors: Evidence for Polyphyletic Origin. Journal of Molecular Evolution 1998, 46, 74–83, doi:10.1007/Pl00006285.

[14] Jiang, C.; Gu, J.; Chopra, S.; Gu, X.; Peterson, T. Ordered origin of the typical two- and three-repeat Myb genes. Gene 2004, 326, 0–22, doi:10.1016/j.gene.2003.09.049.

[15] Liu, C.; Long, J.; Zhu, K.; Liu, L.; Yang, W.; Zhang, H.; Li, L.; Xu, Q.; Deng, X. Characterization of a Citrus R2R3-MYB Transcription Factor that Regulates the Flavonol and Hydroxycinnamic Acid Biosynthesis. Scientific Reports 2016, 6, 25352, doi:10.1038/srep25352.

[16] Phan; A., H.; Li, S.F.; Parish, R.W. MYB80, a regulator of tapetal and pollen development, is functionally conserved in crops. Plant Molecular Biology 2012, 78, 171–183, doi:10.1007/s11103-011-9855-0.

[17] Cheng, H.; Song, S.; Xiao, L.; Soo, H.M.; Cheng, Z.; Xie, D.; Peng, J. Gibberellin acts through jasmonate to control the expression of MYB21, MYB24, and MYB57 to promote stamen filament growth in *Arabidopsis*. PLoS Genet 2009, 5, doi:10.1371/journal.pgen.1000440.

[18] Seo, P.J.; Park, C.M. MYB96-mediated abscisic acid signals induce pathogen resistance response by promoting salicylic acid biosynthesis in *Arabidopsis*. New Phytologist 2010, 186, 471–483. doi:10.1111/j.1469-8137.2010.03183.x.

[19] Zhang, L.; Zhao, G.; Jia, J.; Liu, X.; Kong, X. Molecular characterization of 60 isolated wheat MYB genes and analysis of their expression during abiotic stress. J Exp Bot 2012, 63, 203–214, doi:10.1093/jxb/err264.

[20] Shin, D.H.; Choi, M.G.; Kang, C.S.; Park, C.S.; Choi, S.B.; Park, Y.I. A wheat R2R3-MYB protein PURPLE PLANT1 (TaPL1) functions as a positive regulator of anthocyanin biosynthesis. Biochem Biophys Res Commun. 2016, 469, 686–691, doi:10.1016/j.bbrc.2015.12.001.

[21] Zhang, L; Dong, C; Zhang, Q; Zhao, G; Li, F; Xia, C; Zhang, L; Han, L; Wu, J; Jia, J; Liu, X; Kong, X. The wheat MYB transcription factor TaMYB18 regulates leaf rolling in rice. Biochem Biophys Res Commun. 2016, 481, 77–83, doi:10.1016/j.bbrc.2016.11.014.

[22] Zhang, L.; Zhao, G.; Xia, C.; Jia, J.; Liu, X.; Kong, X. A wheat R2R3-MYB gene, TaMYB30-B, improves drought stress tolerance in transgenic *Arabidopsis*. J Exp Bot 2012, 63, 5873–5885, doi:10.1093/jxb/ers237.

[23] Zhao, Y.; Cheng, X.; Liu, X.; Wu, H.; Bi, H.; Xu, H. The Wheat MYB Transcription Factor TaMYB(31) Is Involved in Drought Stress Responses in *Arabidopsis*. Front Plant Sci 2018, 9, doi:10.3389/fpls.2018.01426.

[24] Shan, T.; Rong, W.; Xu, H.; Du, L.; Liu, X.; Zhang, Z. The wheat R2R3-MYB transcription factor TaRIM1 participates in resistance response against the pathogen Rhizoctonia cerealis infection through regulating defense genes. Sci Rep 2016, 6, 28777, doi:10.1038/srep28777.

[25] Appels, R.; Eversole, K.; Feuillet, C.; Keller, B.; Rogers, J.; Stein, N.; Pozniak, C.J.; Stein, N.; Choulet, F.; Distelfeld, A., et al. Shifting the limits in wheat research and breeding using a fully annotated reference genome. Science (New York, N.Y.) 2018, 361, doi:10.1126/science.aar7191.

[26] Cannon, S.B.; Mitra, A.; Baumgarten, A.; Young, N.D.; May, G. The roles of segmental and tandem gene duplication in the evolution of large gene families in *Arabidopsis thaliana*. BMC Plant Biol 2004, 4, 10, doi:10.1186/1471-2229-4-10.

[27] Xie, T.; Chen, C.; Li, C.; Liu, J.; Liu, C.; He, Y. Genome-wide investigation of WRKY gene family in pineapple: evolution and expression profiles during development and stress. BMC Genomics 2018, 19, 490, doi:10.1186/s12864-018-4880-x.

[28] Panchy, N.; Lehti-Shiu, M.; Shiu, S.H. Evolution of Gene Duplication in Plants. Plant Physiol 2016, 171, 2294–2316, doi:10.1104/pp.16.00523.

[29] Jain, M.; Khurana, P.; Tyagi, A.K.; Khurana, J.P. Genome-wide analysis of intronless genes in rice and *Arabidopsis*. Funct Integr Genomics 2008, 8, 69–78, doi:10.1007/s10142-007-0052-9.

[30] Agarwal, P.; Mitra, M.; Banerjee, S.; Roy, S. MYB4 transcription factor, a member of R2R3-subfamily of MYB domain protein, regulates cadmium tolerance via enhanced protection against oxidative damage and increases expression of PCS1 and MT1C in *Arabidopsis*. Plant Sci 2020, 297, 110501, doi:10.1016/j.plantsci.2020.110501.

[31] Deng, J.; Wu, D.; Shi, J.; Balfour, K.; Wang, H.; Zhu, G.; Liu, Y.; Wang, J.; Zhu, Z. Multiple MYB Activators and Repressors Collaboratively Regulate the Juvenile Red Fading in Leaves of Sweetpotato. Front Plant Sci 2020, 11, 941, doi:10.3389/fpls.2020.00941.

[32] Zhang, J. Evolution by gene duplication: an update. Trends Ecol. Evol. 2003, 18, 292–298, doi:10.1016/S0169-5347(03)00033-8.

[33] Mandaokar, A.; Browse, J. MYB108 acts together with MYB24 to regulate jasmonate-mediated stamen maturation in *Arabidopsis*. Plant Physiol 2009, 149, 851–862, doi:10.1104/pp.108.132597.

[34] Himi, E.; Taketa, S. Barley Ant17, encoding flavanone 3-hydroxylase (F3H), is a promising target locus for attaining anthocyanin/proanthocyanidin-free plants without pleiotropic reduction of grain dormancy. Genome 2015, 58, 43–53, doi:10.1139/gen-2014-0189.

[35] Bi, H.; Luang, S.; Li, Y.; Bazanova, N.; Morran, S.; Song, Z.; Perera, M.A.; Hrmova, M.; Borisjuk, N.; Lopato, S. Identification and characterization of wheat drought-responsive MYB transcription factors involved in the regulation of cuticle biosynthesis. J Exp Bot 2016, 67, 5363–5380, doi:10.1093/jxb/erw298.

[36] Edgar, R.C. MUSCLE: multiple sequence alignment with high accuracy and high throughput. Nucleic Acids Res 2004, 32, 1792–1797, doi:10.1093/nar/gkh340.

[37] Kumar, S.; Stecher, G.; Tamura, K. MEGA7: Molecular Evolutionary Genetics Analysis Version 7.0 for Bigger Datasets. Mol Biol Evol 2016, 33, 1870–1874, doi:10.1093/molbev/msw054.

[38] Wang, Y.; Tang, H.; Debarry, J.D.; Tan, X.; Li, J.; Wang, X.; Lee, T.H.; Jin, H.; Marler, B.; Guo, H., et al. MCScanX: a toolkit for detection and evolutionary analysis of gene synteny and collinearity. Nucleic Acids Res 2012, 40, e49, doi:10.1093/nar/gkr1293.

[39] Krzywinski, M.; Schein, J.; Birol, I.; Connors, J.; Gascoyne, R.; Horsman, D.; Jones, S.J.; Marra, M.A. Circos: an information aesthetic for comparative genomics. Genome research 2009, 19, 1639–1645, doi:10.1101/gr.092759.109.

[40] Yu, G.; Smith, D.; Zhu, H.; Guan, Y.; Lam, T. ggtree: an R package for visualization and annotation of phylogenetic trees with their covariates and other associated data. Methods in Ecology and Evolution 2016, 10.1111/2041-210X.12628, doi:10.1111/2041-210X.12628.

